# Underlying causes for prevalent false positives and false negatives in STARR-seq data

**DOI:** 10.1101/2023.03.03.530915

**Authors:** Pengyu Ni, Siwen Wu, Zhengchang Su

## Abstract

STARR-seq and its variants have been widely used to characterize enhancers. However, it has been reported that up to 87% of STARR peaks are located in repressive chromatins and are not functional in the tested cells. While some of the STARR peaks in repressive chromatins might be active in other cell/tissue types, some others might be false positives. Meanwhile, many active enhancers may not be identified by the current STARR-seq methods. However, the prevalence of and underlying causes for the artifacts are not fully understood. Based on predicted *cis*-regulatory modules (CRMs) and non-CRMs in the human genome as well as predicted active CRMs and non-active CRMs in a few human cell lines with STARR-seq data available, we reveal prevalent false positives and false negatives in STARR peaks and possible underlying causes. Our results will help design strategies to improve STARR-seq methods and interpret the results.

## Introduction

*Cis*-regulatory modules (CRMs) formed by closely located clusters of transcription factor (TF) binding sites (TFBSs) are as important as coding DNA sequences (CDSs) in specifying complex traits of animals (1,2). It is essential to categorize all CRMs and constituent TFBSs in the genomes and determine their functional states and target genes in various cell types in humans and important organisms (3). A CRM often functions independently of its location and orientation relative to the target genes (4). A CRM regulates the transcription of a target gene through interactions between constituent TFBSs in the CRM and cognate TFs in certain cell types where the former are accessible and the latter are available to bind (4,5). Enhancers that increase the expression of target genes are arguably the most important and best-studied type of CRMs. Although more than a million CRMs have been predicted in the human genome, only a few thousand of them have been experimentally validated as enhancers using reporter assays in transgene animal models (6). More recently, various massively parallel assays have been developed to characterize and validate enhancers (7). In particular, self-transcribing assay of regulatory regions sequencing (STARR-seq) originally developed for small genomes clones randomly sheared DNA segments between a minimal-promoter-driven green fluorescence protein open reading frame and a downstream polyA sequence (8). If a sequence is an active enhancer, this results in the transcription of the enhancer sequence, allowing to assess potentially all DNA segments in the genome. STARR-seq and its variants such as WHG-STARR-seq (9) and ATAC-STARR-seq (10), ChIP-STARR-seq (11) and Cap-STARR-seq (12) have been applied to mammal cells, and a great deal of insights into enhancers have been gained from these studies (11,13-16). However, artifacts related to episomal expression vectors used in these methods have been noted (8-11,17-20). On the one hand, it has been shown that 31%, 52% and 87% of STARR peaks identified in fly S2 cells (8), GM12878 cells (10) and LNCaP cells(9), respectively, are not active in their native repressive chromatins. Although the STARR peaks identified in repressive euchromatins of tested cells might be bona fide enhancers that might be active in other cell/tissue types, those found in heterochromatins might be false positives. On the other hand, active enhancers can be missed by the STARR-seq methods due to the length limitation of inserted sequences(20), resulting in false negatives. Nonetheless, the prevalence of such false positives and false negatives and the underlying cause are not fully understood. Based on our recently predicted CRMs and non-CRMs with high accuracy in 85% of the human genome (21) as well as our predicted active CRMs and non-active CRMs (21) in five human cell lines (A549, HCT116, HepG2, K562 and MCF-7) with WHG-STARR-seq data available, and in a human cell line (GM12878) with ATAC-STARR-seq data available, we addressed these important issues.

## Methods The datasets

We downloaded from the PCRMs database (http://cci-bioinfo.uncc.edu) (22) 1,426,947 CRMCs and 1,755,876 non-CRMCs predicted in 85% of the human genome regions using the dePCRM2 pipeline (21,23). We downloaded from the ENCODE data portal (https://www.encodeproject.org/functional-characterization-experiments/) the coordinates of WHG-STARR-seq peaks in A549, HCT116, HepG2, K562 and MCF-7 cell lines. We downloaded the coordinates of ATAC-STARR-seq peaks in the GM12878 cell line from (10). If there were replicates of data in a cell line, we combined them into one set. The dataset is summarized in Supplementary table 1. We downloaded from the ENCODE data portal RNA-seq gene expression data in A549, HCT116, HepG2, K562, MCF-7 and GM12878 cell lines. We downloaded from the Cistrome database(24) the coordinates of epigenetic marks (CA (ATAC-seq), H3K4me1, H3K27ac, H3K4me3) peaks in the six cell lines.

### Prediction of active and non-active CRMs in a cell line

We used our previously trained logistic regression-based universal functional state predictor (UFSP) to predict the functional state (active or non-active) of all the 1,426,947 CRMs in the cell lines using signals of four epigenetic marks CA (ATAC-seq), H3K4me1, H3K4me3 and H3K27ac as previously described (21). Briefly, we trained the UFSP on a negative set made up of all the predicted non-CRMs in the human and mouse genomes and a positive set made by pooling the positive set in each of 67 human and 64 mouse cell/tissue types (21). A positive set in a cell/tissue type is the CRMs that overlap at least a binding peak of TF ChIP-seq datasets collected in the cell/tissue types(21). The signals of the four epigenetic marks CA (ATAC-seq), H3K4me1, H3K4me3 and H3K27ac on the sequences were used as the features.

### Association of sequence elements to genes

As accurate prediction of target genes to distal CRMs is still a highly challenging task, and the existing methods are not necessarily superior to the simple “closest gene assignment” (25), we assigned the gene closest to a sequence element as the target of the latter.

### Construction of heat maps of various chromatin signals

To make the heatmap and density plots, we used all CRMs in the category C regions and all STARR peaks in the categories D and E regions. However, to save computational time, we randomly sampled 10,000 CRMs in the categories A and B regions and the same number of non-CRMs in the category F regions as the STARR peaks in the category E regions. We extracted a 6kb-segment of genome DNA centered on the middle of each sequence, and for each 100-bp sliding window, we calculated the mean of fold enrichment of each chromatin signal (normalized by library size) to input signal (normalized by library size) using EnrichedHeatmap (w0 mode) (26).

### Statistical tests

Two-tailed Mann–Whitney U test was used to compare the gene expression levels, and Kolmogorov– Smirnov (K-S) test was used to compare the distributions of data (density graphs).

## Results

Based on combinatory patterns of TFBSs found in 11,348 TF ChIP-seq datasets for 1,360 TFs in 722 cell/tissue types, whose 1-kbp binding peaks cover 85% of the human genome, we predicted a DNA segment in the genome regions to be either a CRMs or non-CRM, thereby partitioning the genome regions into two exclusive sets: 1.43M CRMs and 1.76M non-CRMs (21,23) (Figure 1A). We predicted each of the CRMs to be either active or non-active in a cell line based on their chromatin accessibility (CA) and three histone marks (H3K4me1, H3K4me3 and H3K27ac) signals (21), thereby partitioning the CRMs into two exclusive sets in the cell/tissue type, i.e., active CRMs and non-active CRMs (Figure 1A). In addition to promoters and enhancers, our predicted CRMs also include silencers and insulators. However, as we used epigenetic marks for active promoter and enhancers to predict the functional states of the CRMs, the predicted active CRMs and non-active CRMs should be mostly active promoters/enhancers and non-active enhancers/promoters, respectively. As promoters may function as enhancers for remote genes(15) and can be detected by STARR-seq methods (8-10), we do not differentiate promoters and enhancers in our analysis, unless specifically pointed out.

**Figure 1.**
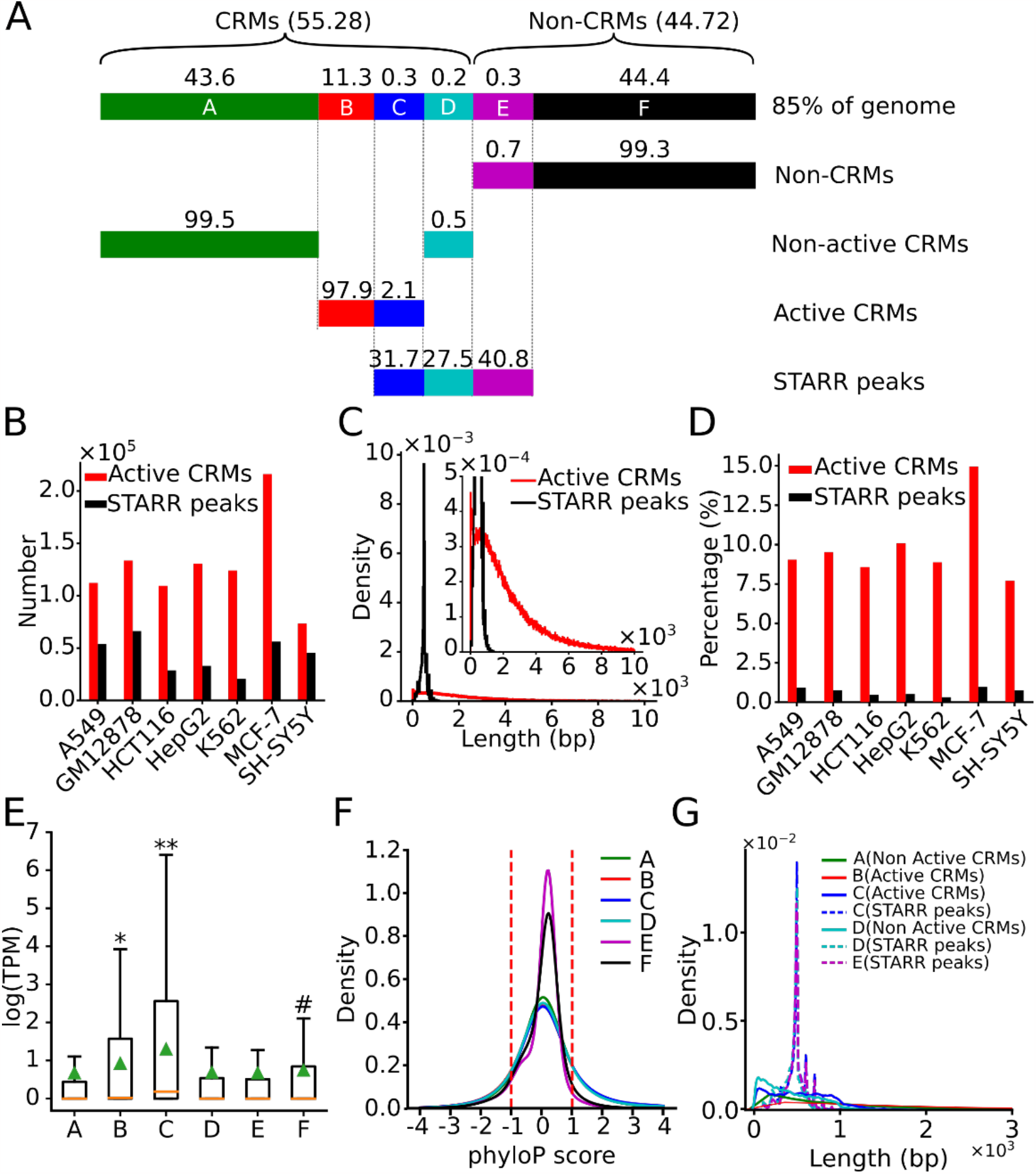
Comparison of our predicted active CRMs and STARR peaks. **A**. Cartoon showing that positions of the 85% of genome regions in a cell line can be exclusively divided into six categories A∼F based on overlaps among predicted active CRMs, predicted non-active CRMs and STARR peaks in a cell line as well as predicted CRMs and predicted non-CRMs in the genome (See text for the definitions of the categories). The number above a line indicates the mean percentage of the category in the indicated category in the six cell lines. For clarity, the lengths of lines are not proportional to the numbers (but see Supplementary Figure 4). **B**. Numbers of predicted active CRMs and STARR peaks in the six cell lines. **C**. Distributions of the lengths of predicted active CRMs and STARR peaks pooled from the six cell lines. **D**. Percentages of genome positions covered by predicted active CRMs and STARR peaks in each of the six cell lines. **E**. Boxplot of expression levels of genes associated with each of the six categories regions; data were pooled from the six cell lines. *p<1.9×10^−4^, comparison between genes associated with the category B regions and those associated with categories A, D, E, and F regions. **p< p<4.90×10^−8^, comparison between genes associated with the category C regions and those associated with other five categories regions. ^#^p=1.36×10^−12^, comparison between genes associated with the category F regions and those associated with the category E regions. All tests were done using two-tailed Mann-Whitney U test. **F**. Distributions of phyloP scores of nucleotide positions of the six categories regions; data were pooled from the six cell lines. **G**. Distributions of the lengths of predicted active CRMs and non-active CRMs in categories regions A∼D, and of STARR peaks in categories regions C∼E; data are pooled from the six cell lines.

### Overlaps between STARR peaks and predicted CRMs and non-CRMs

We predicted at least twice as many active CRMs as the STARR peaks identified in most of the six human cell lines (Figure 1B). Our predicted active CRMs also were much longer than the STARR peaks (median length 1,638 bp vs 489 bp) (Figure 1C and Supplementary Figure 1). This might be due to the size selection step of DNA fragments in constructing expression vector libraries in STARR-seq protocols to cope with the limitation of insertion sizes in expression vectors (20,27). As a result, our predicted active CRMs in each cell line cover a much larger portion of the genome (11.6% on average) than STARR peaks (0.8% on average) (Figure 1D). To see whether we overpredicted active CRMs or STARR-seq methods under determined active CRMs, we analyzed the relation of STARR peaks to our predicted active CRMs, non-active CRMs in a cell line as well as non-CRMs in the 85% of human genome regions. The vast majority (98.8% on average) of genome positions of the STARR peaks in a cell line fell in the 85% of the genome regions, so we ignored the remaining small portion (1.2% on average) that were located in the 15% of the genome regions where we were not able to predict CRMs and non-CRMs due to the lack of TF binding data in these regions (21-23).

Based on overlaps of the four sets of sequences (non-CRMs, non-active CRMs, active CRMs and STARR peaks) in a cell line, we divide the nucleotide positions of the genome regions in the cell line into six categories (A∼F) (Figure 1A): category A consists of nucleotide positions of non-active CRMs that do not overlap the STARR peaks; category B consists of positions of active CRMs that do not overlap the STARR peaks; category C consists of positions of active CRMs that overlap the STARR peaks; category D consists of positions of non-active CRMs that overlap the STARR peaks; category E consists of positions of non-CRMs that overlap the STARR peaks; and category F consists of positions of non-CRMs that do not overlap the STARR peaks. We validated active CRMs, non-CRMs, non-active CRMs and STARR peaks in these category regions using expression levels of the genes that are closest to these elements on linear chromosomes. We say that these closest genes are associated with the relevant category regions.

### Only 31.7% of STARR peaks overlap predicted active CRMs in cell lines

Genes associated with the category C regions (positions of active CRMs that overlap STARR peaks in the cell line) had the highest mean expression levels (p<4.90×10^−8^) among all groups of genes associated with the six categories regions in the pooled six cell lines (Figure 1E), and similar results were seen in each cell line (Supplementary Figure 2). These results suggest that the STARR peaks and their overlapping active CRMs in the category C regions were likely to enhance the expression of the closest genes in their native chromatins. As expected, the category C regions are under strongly evolutionary constraints (Figure 1F and Supplementary Figure 3), suggesting that the STARR peaks and their overlapping predicted active CRMs are at least parts of true CRMs as we argued earlier (21,23). Consistently, CRMs in the category C regions were heavily modified by the active chromatin marks of enhancers H3K27ac, H3K4m1 and H3k4m3, and had high CA as measured by assay for transposase-accessible chromatin using sequencing (ATAC-seq), but were depleted of the repressive enhancer marks H3K9me3 and H3K27me3 (Figure 3 and Supplementary Figure 6). The category C regions comprise an average of 31.7% (C/(C+D+E)) of all STARR peaks positions and 2.1% (C/(B+C)) of all active CRMs positions in the cell lines (Figure 1A and Supplementary Figure 4). Interestingly, the STARR peaks with an average median length of 491bp are slightly shorter than the overlapping active CRMs with an average median length of 521bp in the pooled six cell lines (Figure 1G), and similar results are seen in each cell line (Supplementary Figure 5). Only a small proportion (0.4%∼1.3%) of the STARR peaks include full-length active CRMs, while the majority other STARR peaks contain only parts of overlapping active CRMs (Figure 2A), suggesting that these partial sequences still possess some levels of enhancer activities in STARR-seq assays.

**Figure 2.**
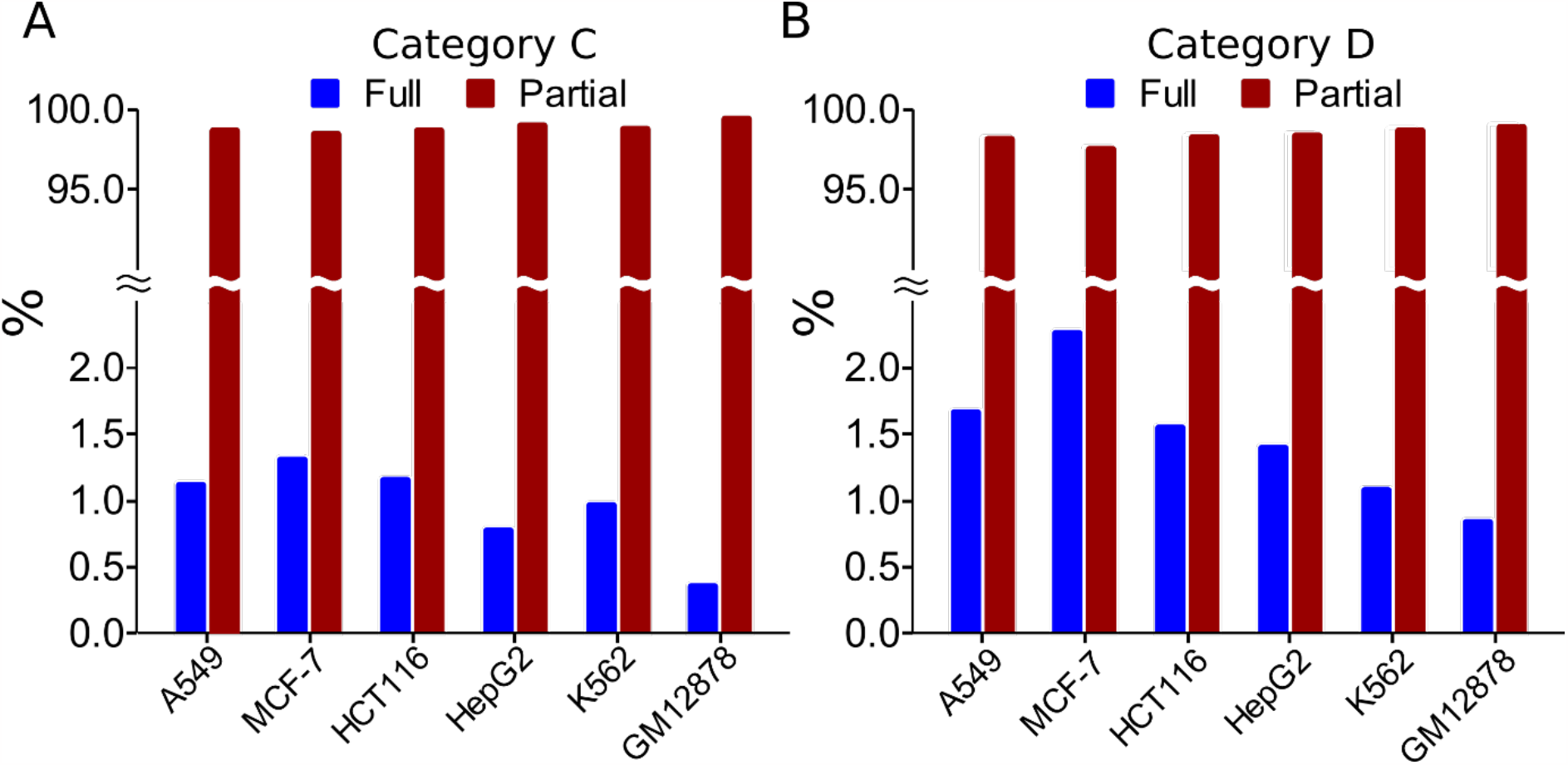
Most STARR peaks in the categories C and D regions include a partial CRM. **A**. Percentages of the STARR peaks containing a full CRM and a partial CRM in the category C regions. **B**. Percentages of the STARR peaks containing a full CRM and a partial CRM in the category D regions.

**Figure 3.**
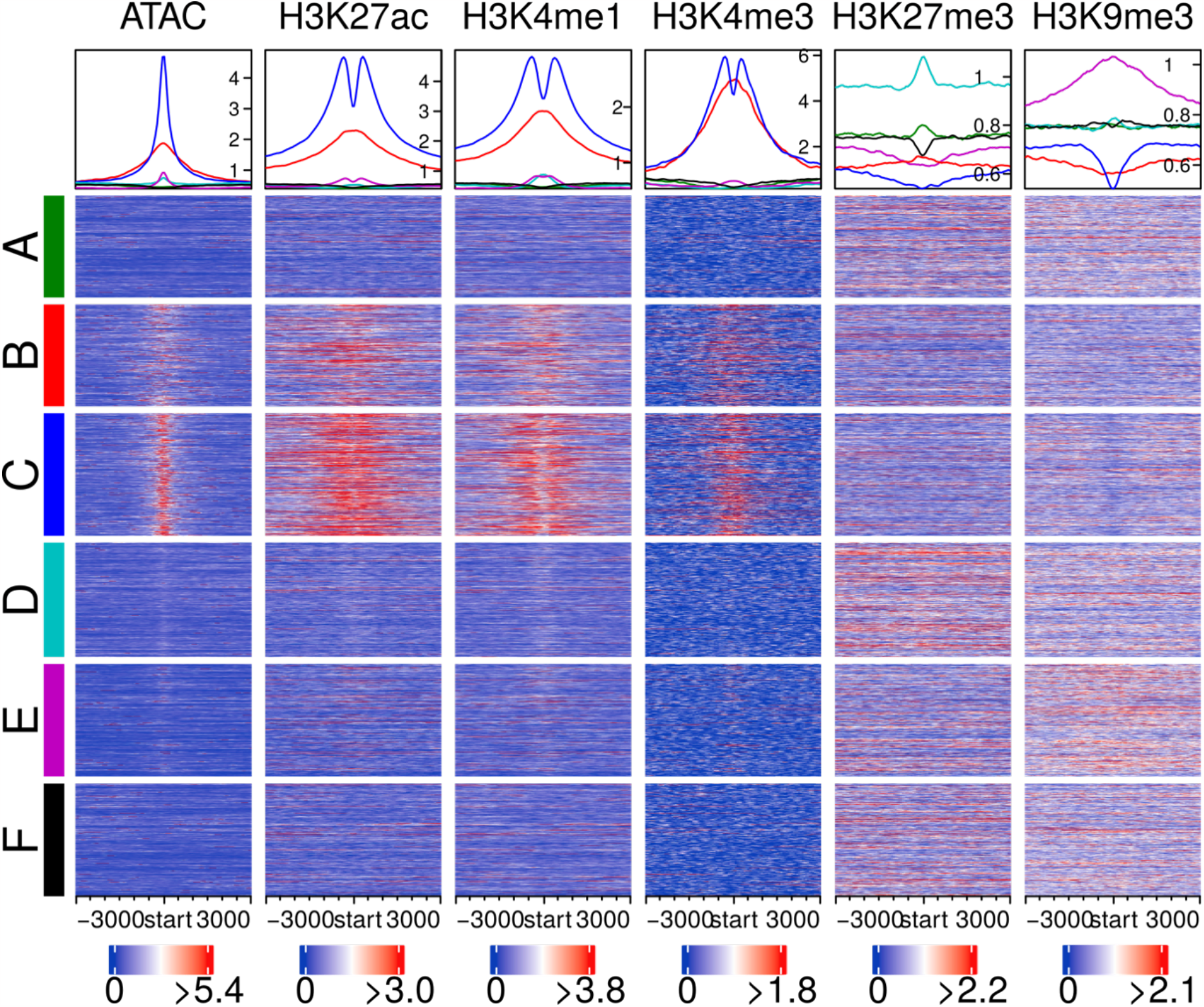
Heat maps of various chromatin signals in the six categories regions in the A549 cells. The heatmap shows the mean signal in each window in each sequence and the density plot shows the mean signal of each window position across all the sequences in the same category regions. The color code for the category regions in the density plot above each column is the same as in the heatmaps.

### Up to 89.5%% of predicted active enhancers might be missed by STARR peaks

Although genes associated with the category B regions (positions of active CRMs that do not overlap STARR peaks in the cell line) had lower expression levels than those associated with the category C regions (p<1.3×10^−10^), their expression levels were significantly higher (p<1.9×10^−4^) than those associated with the other four categories regions (A, D, E and F) in the pooled six cell lines (Figure 1E), and similar results were seen in each cell line (Supplementary Figure 2). These results suggest that the predicted active CRMs in the category B regions were likely to enhance the expression of the closest genes in their native chromatins. As expected, the category B regions are under strongly evolutionary constraints (Figure 1F and Supplementary Figure 3), suggesting that the predicted active CRMs are likely true CRMs as we argued earlier (21,23). Consistently, like CRMs in the category C regions, those in the category B regions were also heavily modified by the active enhancer marks H3K27ac, H3K4m1 and H3K4me3, and had high CA, but were depleted of the repressive enhancer marks H3K9me3 and H3K27me3 (Figure 3 and Supplementary Figure 6).

Interestingly, these predicted active CRMs with an average median length 1,273bp are longer than those in the category C regions with an average median length of 521bp (Figure 1G) in the pooled cells, and similar results are seen in each cell line (Supplementary Figure 5). It is highly likely that the active CRMs in the category B regions might be missed by STARR-seq methods that could only assess sequences shorter than 1,000bp (Figure 1C and Supplementary Figure 1), and the truncated forms of the CRMs that were cloned in the episomal expression vectors might no longer possess enhancer activities. These predicted active CRMs comprise an average of 97.9% (B/(B+C)) of all predicted active CRMs positions in a cell line (Figure 1A and Supplementary Figure 4). Most (89.5%) of these predicted active CRMs are likely distal enhancers as they are least 1,000bp way from the closest downstream transcription start sites (21). These results suggest that STARR-seq methods might have up to an 89.5% false negative rate.

### About 27.5% of STARR peaks are not active in their native repressive euchromatins

Both genes associated with the category A regions (positions of non-active CRMs that do not overlap STARR peaks in the cell line) and genes associated with the category D regions (positions of non-active CRMs that overlap STARR peaks in the cell line) had significantly lower expression levels than genes associated with the categories B and C regions in the pooled cell lines (Figure 1E) and in each cell line (p<7.49×10^−11^) (Supplementary Figure 2), indicating that these predicted non-active CRMs might not enhance the expression of their closest genes in the native chromosomal contexts. As expected, both categories A and D regions are under strongly evolutionary constraints (Figure 1F and Supplementary Figure 3), suggesting that these predicted non-active CRMs are likely to be true CRMs as we argued earlier (21,23), but are in non-active state in their native chromatins.

Consistently, CRMs in both categories A and D regions were depleted of active enhancer marks H3K27ac, H3K4m1 and H3K4me3, and had low CA. However, CRMs in categories A and D regions differ in their repressive enhancer marks in that the former were moderately modified by both the Polycomb-mediated repressive mark H3K27me3 and the heterochromatin repressive mark H3K9me3, while the latter are heavily modified by H3K27me3 but moderately modified by H3K9me3 (Figure 3 and Supplementary Figure 6). The underlying reason for the difference is not clear to us.

Nonetheless, although the STARR peaks in the category D regions might be at least parts of the overlapping non-active CRMs, they are not active in the native chromatins in the tested cell lines. The STARR peaks in the category D regions comprise an average of 27.5% (D/(C+D+E)) of all STRARR peaks positions in the six cell lines, while the overlapping CRMs in category D regions only consist of an average of 0.5% of non-active CRMs positions (Figure 1A and Supplementary Figure 4).

Interestingly, with an average median length of 480bp, the STARR peaks in the category D regions tend to be slightly longer than the overlapping non-active CRMs with an average median length of 351bp in the pooled cell lines (Figure 1G) and in each cell line (Supplementary Figure 5). Still, only a small proportion (0.8%∼2.3%) of the STARR peaks in the category D include a predicted full non-active CRM, while the majority others only contain parts of the overlapping non-active CRMs (Figure 2B). Therefore, non-active CRMs in the category D might become active in episomal expression vectors presumably due to the lack of repressive enhancer marks as has been noted in earlier studies (8,20,27).

### About 40.8% of STARR peaks are enhancer-like sequences masked in heterochromatins

Genes associated with the category E regions (positions of non-CRMs that overlap STARR peaks in the cell line) also had significantly lower expression levels than genes associated with the categories B and C regions (p<1.9×10^−4^), suggesting that the STARR peaks in the category E regions did not enhance the expression of the closest genes in their native chromatins. Interestingly, these STARR peaks were depleted of active enhancer marks H3K27ac, H3K4m1 and H3K4me3, and had low CA, while being heavily modified by the heterochromatin repressive mark H3K9me3 but not by the Polycomb-mediated repressive mark H3K27me3 (Figure 3 and Supplementary Figure 6). These results suggest that the STARR peaks in the category E regions might largely reside in heterochromatins in the tested cell lines. Moreover, the category E regions are more likely evolutionarily neutral than the other four categories (A, B, C and D) (Figure 1F and Supplementary Figure 3, <2.23×10^−302^, K-S test), suggesting that the STARR peaks in the category regions might not be CRMs as we argued earlier (21,23). With an average median length of 492bp, the STARR peaks have similar length distributions to those in the categories C and D (Figure 1G) in the pooled cell lines and in each cell line (Supplementary Figure 5). It is highly likely that the STARR peaks in the category E regions are false positives, which comprise an average of 40.8% (E/(C+D+E) of all STARR peaks positions in the cell lines (Figure 1A and Supplementary Figure 4).

Therefore, the earlier reported STARR peaks in repressive chromatins (8,10,20,28) actually fall into two types: those in the category D regions that are located in repressive euchromatins marked by H3K27me3, and those in the category E regions that are located in repressive heterochromatins marked by H3K9me3. The first type of the STARR peaks in repressive euchromatins might be at least parts of bona fide CRMs that are in poised state in the tested cell line, and they might become active in another cell/tissue types with active chromatin contexts as demonstrated earlier (9,20). The second type in heterochromatins might not be CRM at all, and thus are likely false positives, however, they have not been widely noted before. It follows that these false positive STARR peaks should be reduced by techniques such as ATAC-STARR-seq that depletes heterochromatins before inserting sequences in expression vectors. Indeed, we found that STARR peaks produced by ATAC-STARR-seq in GM12878 cells have a lower proportion (28.9% vs an average of 42.3%) of the second type than those produced by WHG-STARR-seq in the other five cell lines (Supplementary Figure 4). These false positives might arise when enhancer-like sequences that are largely selectively neutral (Figure 1F and Supplementary Figure 3) and mostly masked in their native heterochromatins (Figure 3 and Supplementary Figure 6) become active in episomal expression vectors. The two types of STARR peaks in repressive chromatins together comprise 53.3% of ATAC-STARR peaks positions in GM12878 cells and an average of 68.3% of WHG-STARR peaks positions in the other five cell lines (Supplementary Figure 4). Remarkably, the total proportion of STARR peaks that we inferred to be in either repressive euchromatins (24.5%) or heterochromatins (28.9) in GM12878 cells (53.4%) (Supplementary Figure 4) is in good agreement with the earlier estimate of 52% in this cell line(10).

### Enhancer-like sequences masked in heterochromatins might be evolved by genetic drift

Notably, genes associated with the category F regions (positions of non-CRMs that do not overlap STARR peaks in the cell line) also had significantly lower expression levels than those associated with the categories B and C regions (p<4.90×10^−8^), but significantly higher expression levels than those associated with the category E regions (p=1.36×10^−12^) in pooled cells and in four out of the six cell lines (p<3.17×10^−4^) (Figure 1E and Supplementary Figure 2). Non-CRMs in the category F regions were depleted of the active enhancer marks H3K27ac, H3K4m1 and H3K4me3, and had low CA, but were moderately modified by the repressive enhancer marks H3K9me3 and H3K27me3 (Figure 3 and Supplementary Figure 6). As expected, like the category E regions, the category F regions are also more likely evolutionarily neutral than the other four categories (A, B, C and D) (p<2.23×10^−302^, K-S test), suggesting that the non-CRMs in the category F regions are not CRMs as we argued earlier (21,23). Interestingly, the category E regions are more likely to be evolutionarily neutral than the category F regions (Figure 1F and Supplementary Figure 3). The possible reason might be that less evolutionary constraints in heterochromatins (29) render the STARR peaks in the category E regions to become CRM-like sequences by genetic drift without causing deleterious effects as they are masked in heterochromatins, while unwanted CRM-like sequences in euchromatin can be deleterious, and are subject to purifying selection (30). This also explain why STARR peaks in the category E regions show enhancer functions in episomal expression vectors. The fact that the category E regions are heavily modified by the heterochromatin mark H3k9me3, but the category F regions are only moderately modified by H3k9me3 (Figure 3 and Supplementary Figure 6), might at least partially explain why genes closest to the former category have lower expression levels in native chromatins than those closest to the latter category (Figure 1E and Supplementary Figure 2).

## Discussion

In this study, we identified two types of STARR peaks that are located in repressive chromatins in the tested cell lines but display enhancer activity in episomal expression vectors. The first type makes up an average of 27.5% of WHG-STARR peaks and 24.5% of ATAC-STARR peaks. They overlap our predicted non-active CRMs that are depleted of active enhancer marks but heavily modified by the Polycomb-mediated repressive mark H3K27me3 and moderately modified by the heterochromatin repressive mark H3K9me3. It is likely that this type of STARR peaks are at least parts of CRMs that might be active in other cell/tissue types. Therefore, this type of STARR peaks is still valuable for characterizing all the regulatory elements encoded in the genome, although they are not active in the tested cell lines. The second type comprises 42.3% of WHG-STARR peaks and 28.9% of ATAC-STARR peaks. They overlap our predicted non-CRMs, and thus are likely false positives. These STARR peaks are depleted of active enhancer marks and heavily modified by the heterochromatin repressive mark H3K9me3 but not by the Polycomb-mediated repressive mark H3K27me3. Thus, they are likely short CRM-like sequences masked in heterochromatins. As this type of STARR peaks might be false positives, therefore, ideally, they should be maximally reduced in the STARR peaks.

Our findings suggest a strategy to reduce the second type of STARR peaks that are located in heterochromatins. For example, by enriching naked DNA using ATAC when preparing insertion sequences from genomic DNA, thereby depleting heterochromatins, ATAC-STARR-seq (10) has a much lower proportion (28.9%) of STARR peaks in heterochromatin than does WHG-STARR-seq (9) (42.3%) that does not enrich naked DNA. A more efficient method for depleting heterochromatins during insertion sequence preparation is needed to further reduce the proportion of the second type of STARR peaks in heterochromatins. One possible method is to combine a ATCA or DNase digestion-based method with a ChIP step targeting marks such as H3K4me1 or H3K27ac for active and poised enhancers.

We also found that 97.9% of genome positions of our predicted active CRMs in a cell line were missed by the STARR peaks, and up to 89.5% of these predicted active CRMs are likely distal enhancers (21). In other words, current STARR-seq methods with a false negative rate up to 89.5%, are only able to detect less than 10.5% of active CRMs in a cell line. Since most known enhancers are longer than 500bp (for instances, enhancers in the VISTA database have a mean length o f2,049bp (21)), and our predicted CRMs have a mean length about 1,000bp, while the plasmid (9,10) and lentivirus (18) used in current STARR-seq methods accommodate an insertion size up to 500bp. Therefore, it is highly likely that these active enhancers cannot be detected by the STARR-seq methods because they are too long to be inserted in episomal expression vectors, while their truncated short insertable forms might not be functional. Our findings suggest a strategy to reduce the false negative rate. For example, using expression vectors that can accommodate longer insertion sequences might allow to detect longer enhancers. However, some very long enhancers (with a length of several thousand bp) cannot be inserted in most known expression vectors, and therefore different methods such as CRISPR interference (31,32) might be needed to validate them.

## Data availability

The authors declare that all data supporting the findings of this study are available in the paper and its supplementary information files.

## Acknowledgements

We would like to thank all lab members for their discussion. The work was supported by US National Science Foundation (DBI-1661332). The funding bodies played no role in the design of the study and collection, analysis, and interpretation of data and in writing the manuscript.

## Author contributions

ZS and PN conceived the project. ZS and PN developed the algorithms. PN and SW carried out computational experiments and analysis. PN and ZS wrote the manuscripts. All authors read and approved the final manuscript.

## Competing interests

The authors declare no competing financial interests.

## Supplementary information

Supplementary information is available for this paper online.

## Notes

### Competing Interest Statement

The authors have declared no competing interest.

