## Supplementary Figures for "Underlying causes for prevalent false positives and false negatives in STARR-seq data"

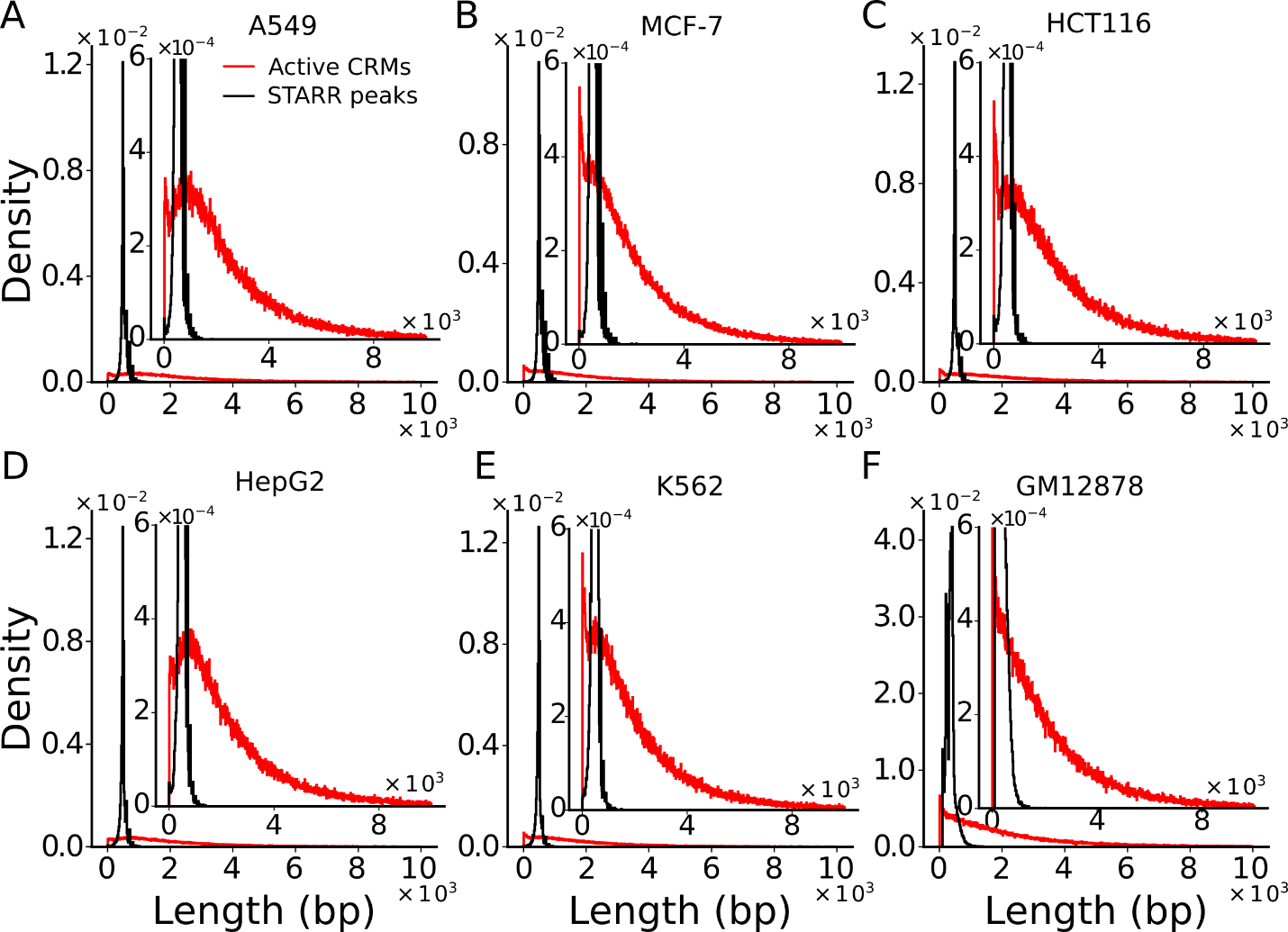

**Supplementary Figure 1. Distributions of the lengths of predicted active CRMs and STARR peaks in each cell line.** The insets are blow-up views of the indicated in regions of the axes.

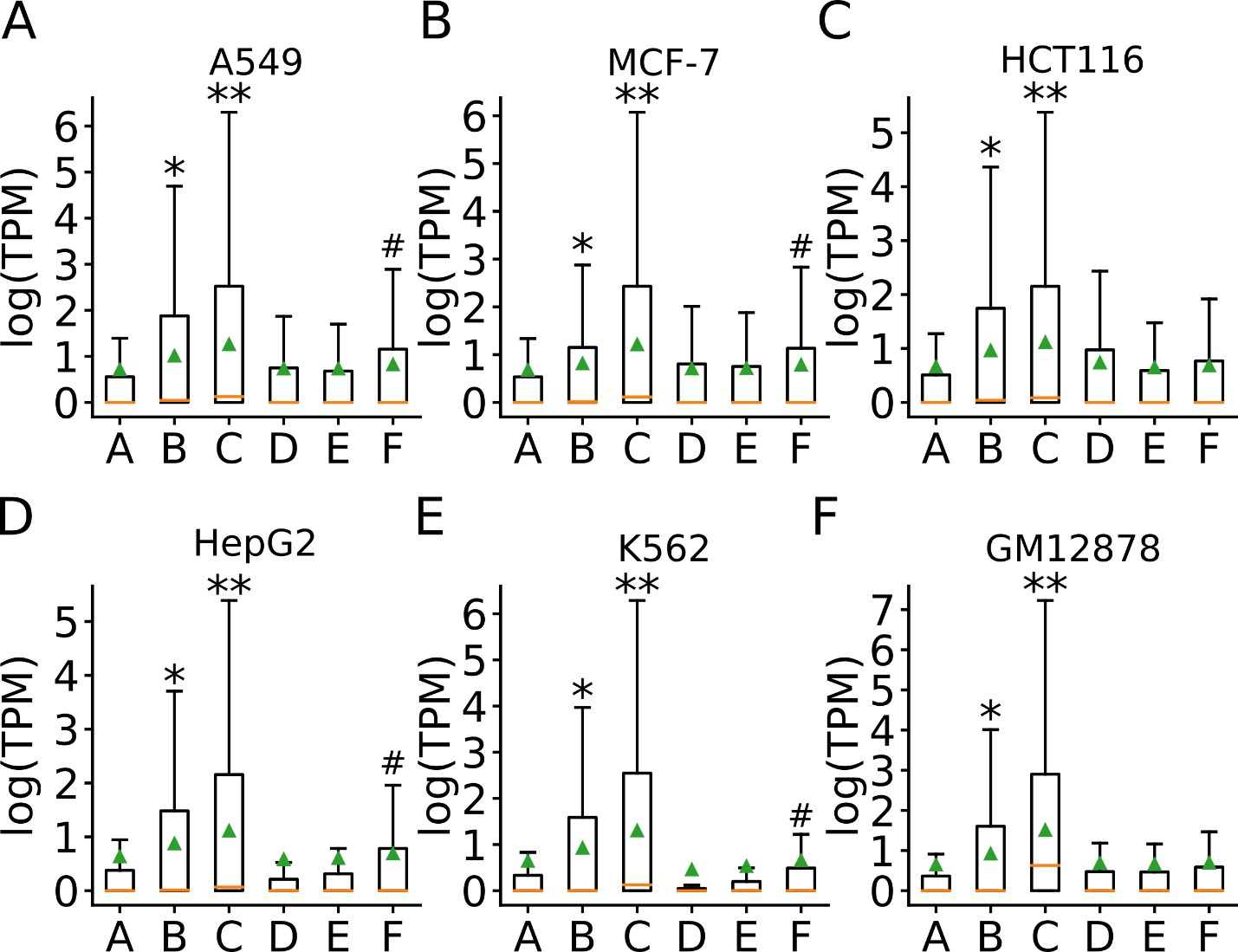

**Supplementary Figure 2. Expression levels of closest genes to the predicted active CRMs, non-active CRM and non-CRMs as well as STARR peaks in the six categories regions in each cell line.** **p<0.001, comparison between genes associated with the category B regions and those associated with categories A, D, E, and F regions. **p<0.001, comparison between genes associated with the category C regions and those associated with other five categories regions. ^#^p<0.001, comparison between genes associated with the category F regions and those associated with the category E regions. All tests were done using two-tailed Mann-Whitney U test.

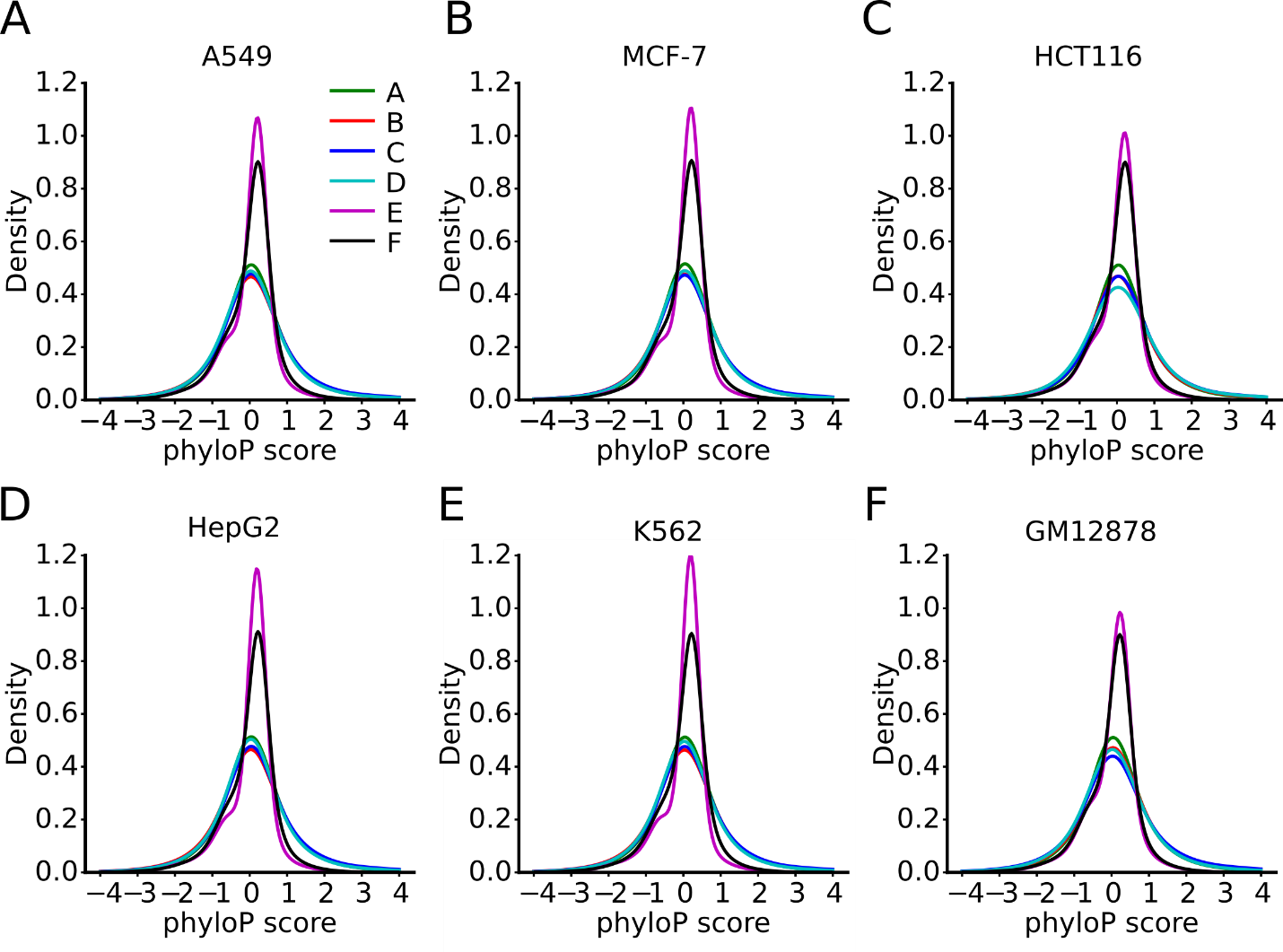

**Supplementary Figure 3. Distributions of phyloP scores of nucleotide positions of the six category regions in each cell line.**

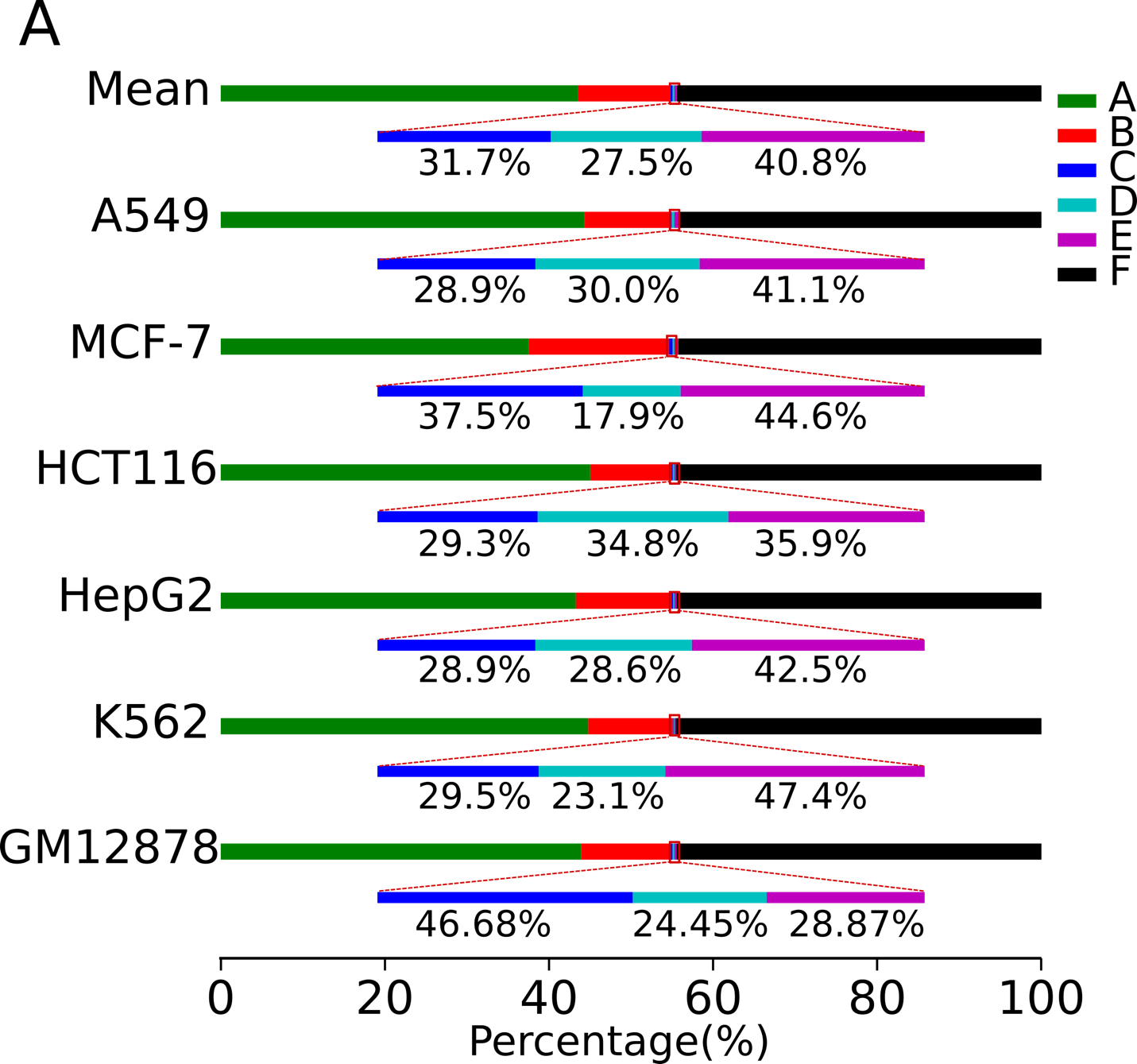

**Supplementary Figure 4. Cartoon showing the percentages of sizes of the six category regions in the genome in each cell line, as well as their means.** The blow-up views show categories C, D and E regions, and their relative proportions. The axis at the bottom indicates the proportion of each category in the genome.

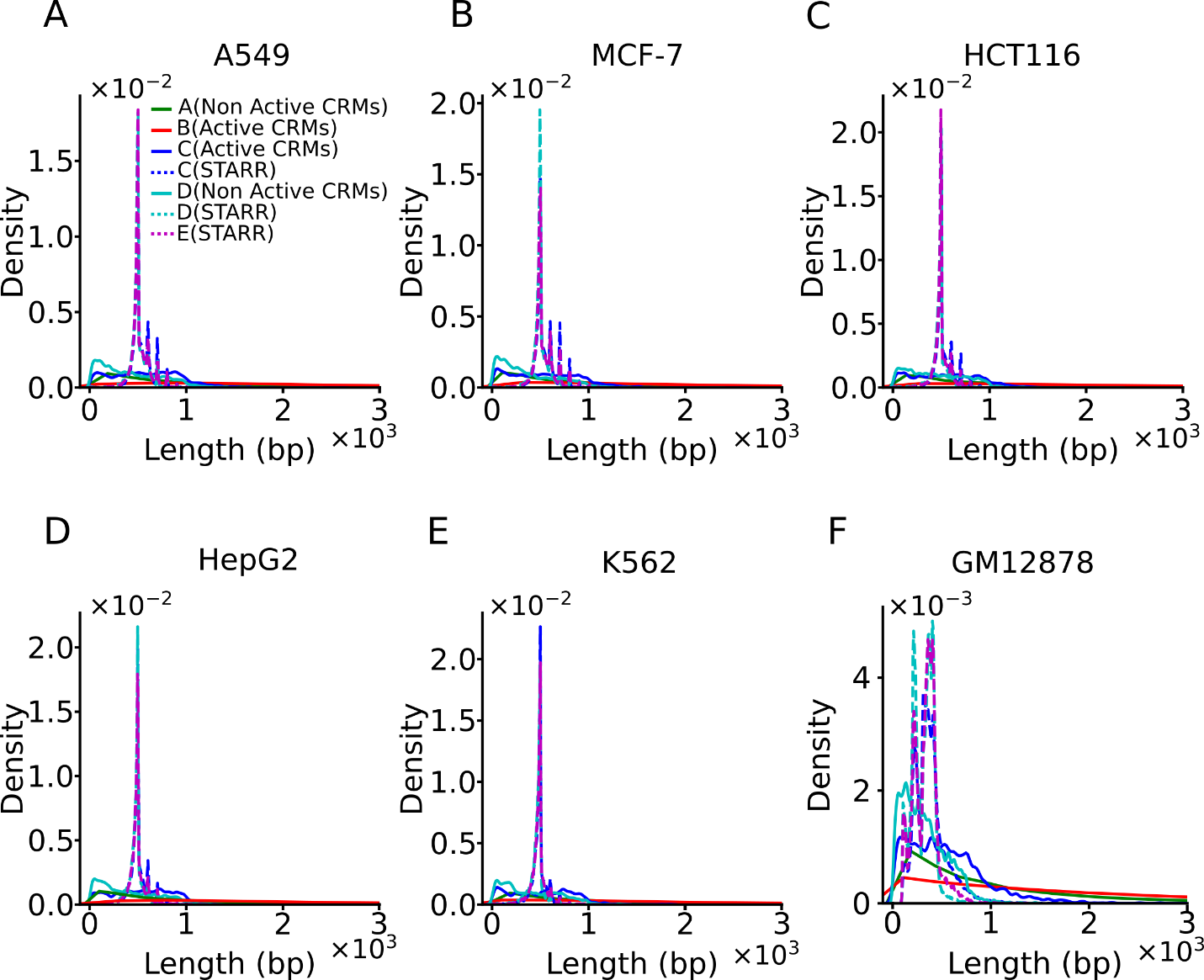

**Supplementary Figure 5. Distributions of the lengths of predicted active CRMs and non-active CRMs in categories A~D regions, and of STARR peaks in categories C~E regions in each of the six cell lines.**

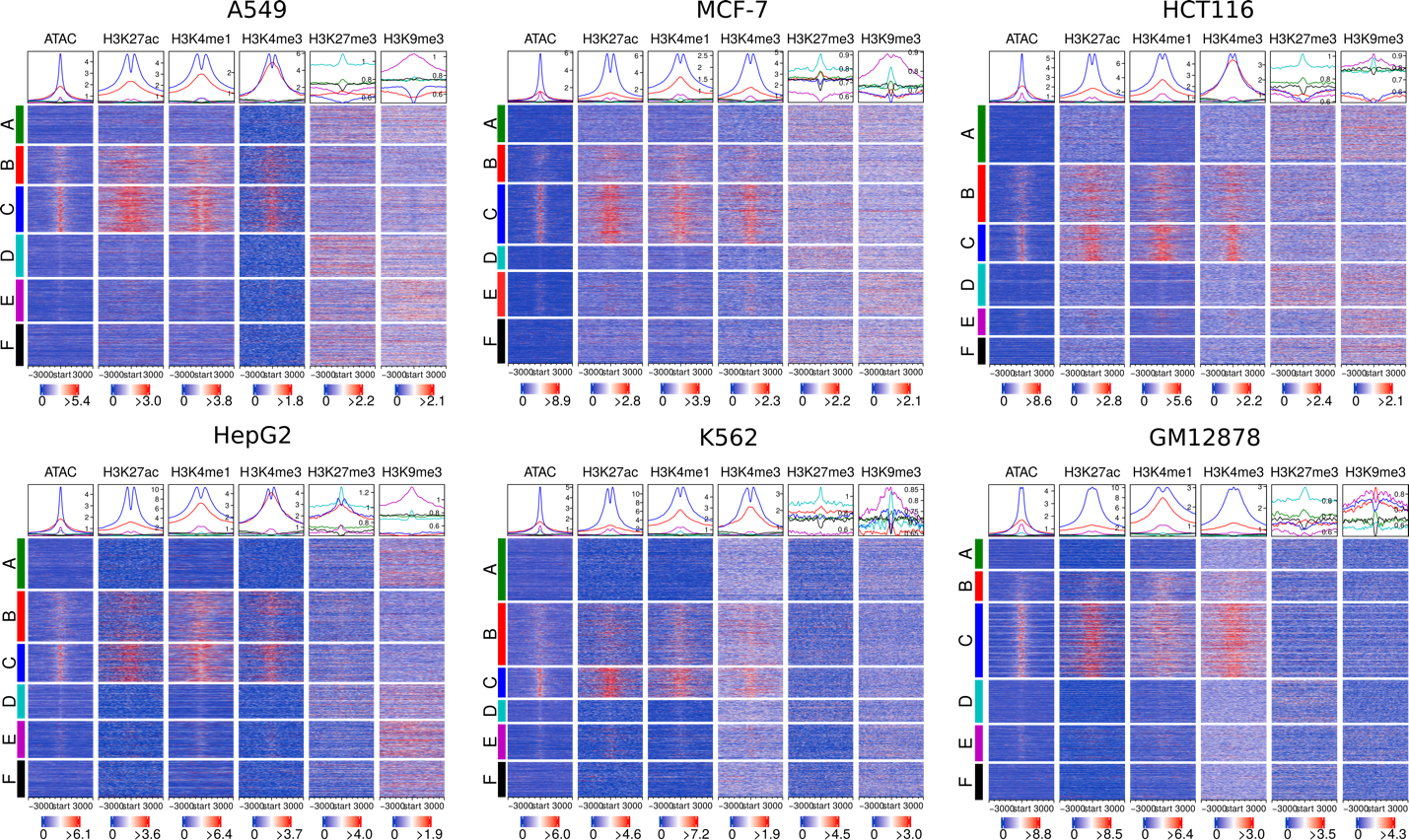

**Supplementary Figure 6. Heat maps of various chromatin signals in the six categories regions in the six cell lines.** The heatmap shows the mean in each window in each sequence and the density plot shows the mean of each window position across all the sequences in the same category regions. The color code for the category regions in the density plot above each column is the same as in the heatmaps.
